# Towards a two-stage model of action-stopping: Attentional capture explains motor inhibition during early stop-signal processing

**DOI:** 10.1101/2021.02.26.433098

**Authors:** Joshua R. Tatz, Cheol Soh, Jan R. Wessel

**Affiliations:** Department of Psychological and Brain Sciences, University of Iowa, Iowa City, IA 52245; Department of Neurology, University of Iowa Hospitals and Clinics, Iowa City, IA 52242

## Abstract

The ability to stop an already initiated action is paramount to adaptive behavior. Most scientific debate in the field of human action-stopping currently focuses on two interrelated questions. First: Which mental and neural processes underpin the implementation of inhibitory control, and which reflect the attentional detection of salient stop-signals instead? Second: Why do physiological signatures of inhibition occur at two different latencies after stop-signals (for visual signals, either before or after ∼150ms)? Here, we address both questions via two pre-registered experiments that combined transcranial magnetic stimulation, electromyography, and multi-variate pattern analysis of whole-scalp electroencephalography. Using a stop-signal task that also contained a second type of salient signal that did not require stopping, we found that both signals induced equal amounts of early-latency inhibitory activity, whereas only later signatures (after 175ms) distinguished the two. These findings resolve ongoing debates in the literature and strongly suggest a two-step model of action-stopping.

## Introduction

One of the core cognitive control abilities available to healthy adult humans is action-stopping: being able to cancel an impending movement even after its initiation. Basic neuroscientific research on this ability, most prominently conducted using the stop-signal paradigm (Logan et al., 1984; Verbruggen et al., 2019) has identified a three-pronged neuroanatomical network – consisting of the right inferior frontal cortex (rIFC), the pre-supplementary motor area (pre-SMA), and the subthalamic nucleus (STN) – which implements an inhibitory control mechanism that purportedly underlies the ability to rapidly stop an action (Aron et al., 2007). However, recent years have been marred by substantial controversy regarding the purported role of this network in uniquely instantiating inhibitory control. In particular, a set of studies using functional magnetic resonance imaging (fMRI) has shown that infrequent events that do not instruct the subject to stop an action produce activity in overlapping brain regions (Hampshire et al., 2010), with some going as far as claiming that “IFC sub-regions are not functionally unique in their sensitivities to inhibitory cognitive demands” (Erika-Florence et al., 2014) and instead reflect the processing of infrequent events. The implication is that rIFC activity after stop-signals is not due to the implementation of inhibitory control, but due to the infrequent nature of the stop-signal.

However, it is also known that the success of stopping, particularly in the stop-signal task (SST), depends strongly on the relative timing of processes related to movement initiation, execution, and cancellation (Logan et al., 1984). Therefore, fMRI notably lacks sufficient temporal resolution to fully resolve debates that pertain to the exact cascade of neural processes underlying action-stopping. Inhibitory control and attentional capture are tightly intertwined (Wessel & Aron, 2017), and it is widely accepted that both are necessary to successfully stop an action in tasks like the SST (Matzke et al., 2013; Elchlepp et al., 2016). To identify which neural signatures and regions relate to either process, methods with better temporal resolution than fMRI are necessary. However, the exact debate outlined above – which stop-related signatures reflect inhibitory control and which are related to the attentional detection of the stop-signal – is currently also taking place in work that uses such methods. For example, studies employing electroencephalography (EEG) have long proposed fronto-central activity such as the P3 event-related potential (ERP) after stop-signals to index inhibitory control (de Jong et al., 1990; Kok et al., 2004; Wessel & Aron, 2015). Several groups, however, have claimed that the P3 reflects a process unrelated to the actual implementation of inhibitory control and have suggested that earlier ERPs may index that function instead (e.g., Huster et al., 2020; Skippen et al., 2020). In the same respect, it is also notable that motor evoked potentials, a measurement of net-corticospinal excitability (CSE) derived from a combination of electromyography (EMG) and transcranial magnetic stimulation (TMS), show that CSE is already broadly suppressed at around 150ms following stop-signals – i.e., before the emergence of such classic purported inhibitory signatures as the P3 ERP (e.g., Majid et al., 2013; Wessel et al., 2013; Jana et al., 2020). This is further corroborated by EMG activity itself: residual subthreshold activation of the target muscle that is observable on many successfully cancelled actions undergoes a visible reduction right around that same latency (Raud & Huster, 2017; Raud et al., 2020). These findings have led to the proposal that the true neural measure of inhibitory control – termed “CancelTime” – is orders of magnitude shorter than previously thought (Jana et al., 2020), and invalidates many later signatures, including the fronto-central P3 and others, as non-critical for inhibitory control (e.g., Hanes et al., 1998; Ogasawara et al., 2018). Hence, instead of resolving the debate from the fMRI literature, research with temporally precise methods further complicated the debate on which neural processes reflect inhibitory control during action-stopping.

Additionally, recent research on the presence of reactive, automatic inhibitory control following salient, surprising, or merely infrequent signals adds another layer of nuance to this discussion. Both EEG (Waller et al., 2019; Novembre et al., 2018; 2019) and CSE studies (Wessel & Aron, 2013; Dutra et al., 2018; Iacullo et al., 2020) have consistently shown that even without an instruction to stop an ongoing or impending action, such salient, surprising, or merely infrequent events automatically induce purportedly inhibitory signatures such as non-selective CSE suppression. This clearly raises the question of which purported inhibitory signatures are uniquely related to outright action-stopping, and which are observable after any type of salient event.

Here, we aimed to resolve these questions using two experiments that measured all three purported signatures of inhibitory control that have sufficient temporal resolution to potentially disentangle infrequent/salient event detection inhibitory control: the non-selective suppression of CSE (Badry et al., 2009), the short-latency suppression of target-muscle EMG on successful stop-trials (Raud & Huster, 2017), and event-related neural activity recorded from the whole scalp EEG, which was subjected to multi-variate pattern analysis (MVPA) to distinguish activity on infrequent/salient events from stop-signals with millisecond resolution. In both experiments, participants performed a version of the SST that included a third signal (in addition to the go- and stop-signals), which was perceptually comparable to the stop-signal, but did not instruct stopping (a logic similar to previous studies such as Hampshire et al., 2010; Dodds et al., 2010; Chatham et al., 2012; and Erika-Florence et al., 2014). In this pre-registered study, we hypothesized that both these no-stop/ignore signals and stop-signals themselves would trigger both low-latency signatures of inhibition at the level of the motor system – i.e., suppression of both CSE and EMG. Moreover, we hypothesized that there would be no early-latency difference in magnitude between these signatures on stop-compared to ignore-signals. We furthermore hypothesized that EEG-MVPA would concomitantly show that stop- and ignore-signals do not result in decodable differences in brain activity until after the initial ∼150ms post-signal period that contains the CSE/EMG suppression. Lastly, we hypothesized that EEG signatures that uniquely decode stop-from ignore-signals would emerge after that initial period.

## Experiment 1

### Method

#### Hypothesis

The pre-registered hypothesis for Experiment 1 was that both salient/infrequent events that do not instruct stopping would induce short-latency CSE suppression, and that that degree of that initial CSE suppression would not differ significantly from that observed following stop-signals. The pre-registration document can be found at https://osf.io/b72mp/?view_only=75676ea4492a41a59df79528e6f81b41.

#### Participants

Twenty-nine healthy, right-handed adults (16 female, mean age = 21.83, *SD* = 4.46) participated in the TMS version of the experiment after completing screening questions (Rossi et al., 2011) and providing written informed consent. The participants were recruited via an email sent out to the University of Iowa community, an online system providing research experience to students in psychology classes, or among personnel in psychology labs. Participants were compensated at an hourly rate of $15 or received course credit. The study was approved by the University of Iowa’s Institutional Review Board (#201711750).

The sample size was determined by an a priori analysis computed in G*Power (Faul et al., 2009) for repeated measures ANOVAs for 1 group and 3 measurements. Based on a recent study that showed non-selective CSE suppression in following infrequent events (Iacullo et al., 2020), we sought 80% power to identity a medium effect (*f* = .25) using a pre-defined alpha level of .05 (other G*power parameters were left to default). Two additional participants were collected to replace participants with insufficient trial counts.

#### Materials, Design, and Procedure

The behavioral task was presented using Psychtoolbox (Brainard, 1997) and MATLAB 2015b (TheMathWorks, Natick, MA) scripts on a Linux desktop computer running Ubuntu. Figure 1 (*top panel*) depicts the behavioral task. Each trial began with a white fixation cross on a black background. After 300 ms, the fixation cross was replaced by a centrally displayed white arrow pointing to the left or right. This represented the GO signal. Participants responded to the direction of the GO signal by pressing the corresponding left or right foot pedal on a Kinesis Savant Elite 2 response device. On 2/3 of trials, termed GO trials, the arrow remained white for 1 s. On 1/6 of trials, the white arrow turned magenta after an adaptive delay (see below). On another 1/6 of trials, the arrow turned from white to cyan after a matched delay. These signals represented the STOP and IGNORE signals, respectively (the color assignment was counterbalanced across subjects). Participants were instructed to try to cancel their response to the STOP signal and to continue responding in the case of an IGNORE signal. Participants were instructed to respond according to the direction of the GO signal and had a maximum deadline of 1 s to do so. If no response was made, the text “Too slow!” was displayed centrally in red for 1 s. A stop was considered successful if, following the STOP signal, no response was registered within the 1 s response deadline. No immediate feedback was provided following STOP signals nor was immediate feedback provided following GO and IGNORE signals (except on misses – i.e., when no response was made on either of these trials types within 1 s). The delay of the color change, known as the stop-signal delay (SSD) in the case of the STOP signal, was initialized at 200 ms and from there adjusted adaptively based on stopping success. If the participant was successful in withholding a response following the STOP signal, the SSD was increased by 50 ms for subsequent trials. If the participant made a response after presentation of the STOP signal (but within the response deadline), the SSD decreased by 50 ms with the constraint that the SSD could not fall below 50 ms. The delay for the IGNORE signal was inherited from the most recent STOP signal. Immediately following the 1 s response deadline, the arrow was again replaced with the fixation cross. The cross was presented for 1.7 s plus a variable inter-trial interval ranging from 150 to 250 ms (sampled from a uniform distribution in 25 ms increments).

**Figure 1.**
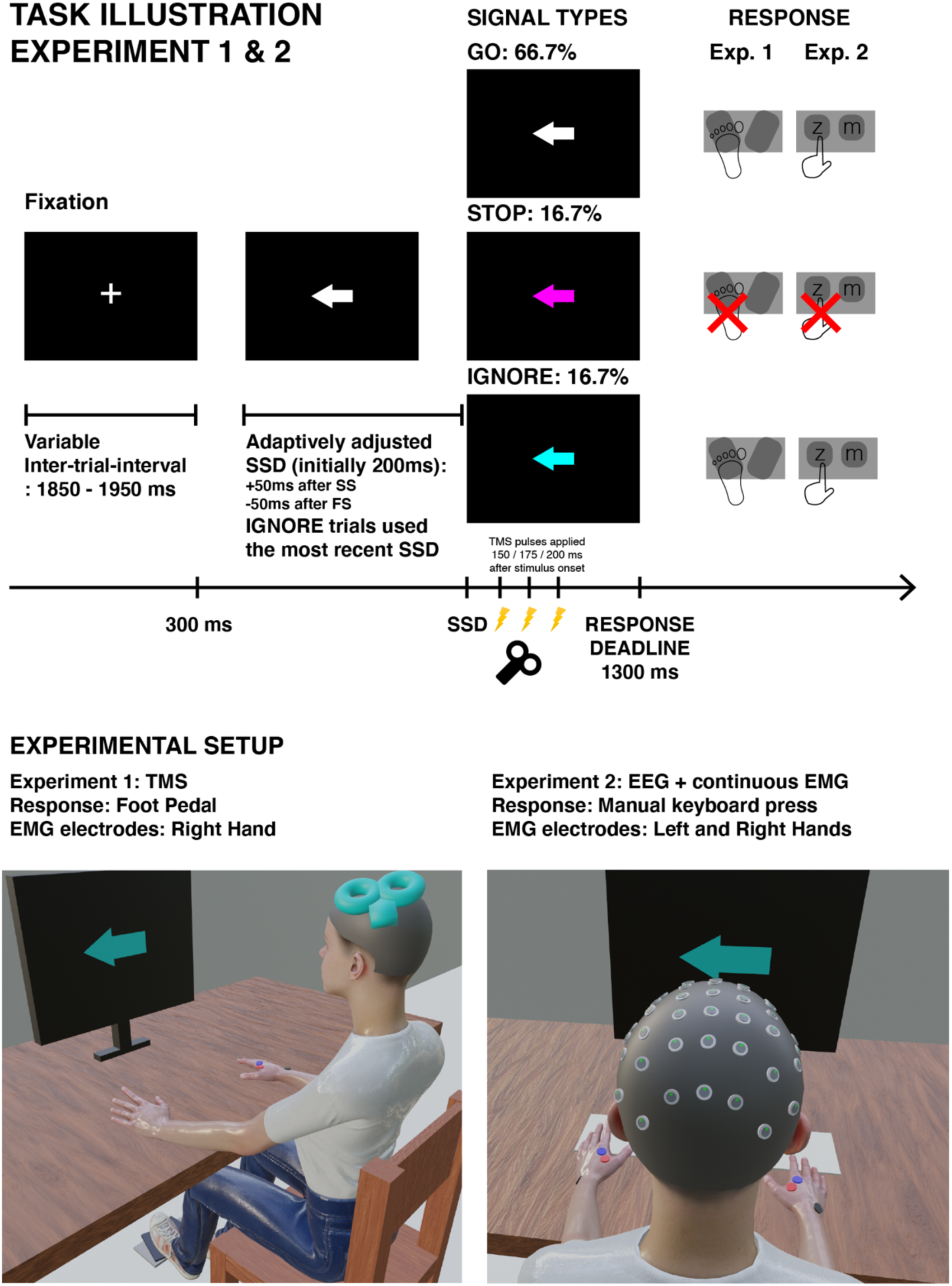
Task Diagram. (Top panel) Illustrates the time course of the behavioral task. Following presentation of the GO stimulus, the arrow changed to magenta or cyan on a minority of trials (16.7% for each color). These signaled to the participant to try to cancel their response (STOP signal) or to – as with the GO signal – respond as quickly as possible in the indicated direction (IGNORE signal). Thus, the STOP and IGNORE signals were similar in all respects but action requirements. (Lower left) The set-up for Experiment 1 which involved recording MEPs from the right hand while the feet were involved in the task. (Lower right) The set-up for Experiment 2 which recorded EEG from the scalp and EMG from both hands which were involved in the task.

Prior to the main experiment, each participant completed a practice block of 36 trials. No TMS was delivered during the practice block, and these data were not included in the analyses. The main experiment featured a 3 (Signal Type: GO, IGNORE, STOP) x 3 (Stimulation Time: 150, 175, 200ms) design with both Signal Type and Stimulation Time as within-subject factors. A random permutation function was used to achieve a pseudorandomized trial order with respect to Signal Type and Stimulation Time. The constraints were that the first 3 trials were necessarily GO trials and that no two Infrequent trials (IGNORE or STOP) were presented consecutively. The arrow direction was randomly determined for each trial (sampling from a Bernoulli distribution). Participants completed 1080 trials in total. Single-pulse TMS was delivered on 840 of those trials, with 180 pulses corresponding to STOP trials, 180 to IGNORE trials, and 240 to GO trials. TMS was delivered 150, 175, or 200 ms post-stimulus onset in equal numbers for each trial type. Thus, TMS was delivered on each STOP and IGNORE trial. Another 240 trials were dedicated to obtaining active baseline MEP measurements, during which, TMS was delivered prior to the onset of the GO signal on GO trials. The remaining 240 trials (without TMS) were all GO trials that preceded the active baseline GO trials. These trials did not have TMS to allow adequate time for the TMS stimulator to recharge between pulses. Active baseline and no TMS trials were selected randomly from the list of all consecutive GO trials (with the constraint that equal numbers of 150, 175, and 200 ms stimulation times were preserved for regular GO trials). Due to an initial programming error, the first participant received 672 GO trials and only 24 active baseline trials (along with the planned 180 STOP trials and 180 IGNORE trials that everyone received). As this participants’ data were usable and representative in all other respects, their data were not excluded from the analyses.

The 1080 trials were partitioned into 10 blocks of 108 trials each. In between blocks, participants were given as much rest time as they desired. In addition, the RT, miss rate, and direction errors were displayed to the participant. If necessary, participants were given instructions based on these metrics as well as p(inhibit) and SSD (which were hidden in the decimal places of the other metrics to inform the experimenter without informing the subject). As typical for the SST, the experimenter’s instructions aimed to keep the participant around p(inhibit) = .5 by encouraging the participant to either respond more quickly or to be more successful at stopping depending on these indicators. In a few blocks (one out of the ten blocks in seven of the 27 participants and two out of the ten blocks in one additional participant), the SSD became longer than 500 ms. When this occurred, the next block was started with the SSD reinitialized to 200 ms, and the experimenter reiterated the importance of responding quickly. Blocks in which p(inhibit) was 0 or 1 or in which the miss (no response following the GO signal) rate was > 25% were excluded from all analyses. This was the case for 3 blocks in total, each from different participants.

#### Motor evoked potentials

We measured motor-evoked potentials (MEPs) as an index of CSE. As we were specifically interested in the ‘global’, non-selective CSE suppression that is typical for reactive inhibition in the stop-signal task (Badry et al., 2009), we measured CSE from a motor effector that was not involved in the task itself. We recorded electromyography (EMG) from the right first-dorsal interosseous (FDI) muscle and stimulated the corresponding region of contralateral M1 using TMS. Participants were instructed to keep both hands relaxed and pronated on the desk while data were collected and they were responding to the task stimuli with their feet.

TMS was delivered via a MagStim 200-2 system (MagStim, Whitland, UK) with a 70-mm figure-of-eight coil. Hotspotting was used to identify the correct location and intensity for each participant. We initially positioned the coil 5 cm left of and 2 cm anterior to the vertex and incrementally adjusted the location and intensity to determine the participant’s resting motor threshold (RMT). RMT was the minimum intensity needed to produce MEPs greater than .1 mV in 5 out of 10 consecutive probes (Rossini et al., 1994). For the main experiment, the stimulus intensity was increased to 115% of RMT. The mean experimental intensity was 58.6% of maximum stimulator output.

EMG was recorded using adhesive electrodes (H124SG, Covidien Ltd., Dublin, Ireland) placed on the belly and tendon of the right FDI. A ground electrode was placed on the distal end of the ulna. EMG electrodes were passed through a Grass P511 amplifier (Grass Products, West Warwick, RI) and sampled using a CED Micro 1401-3 sampler (Cambridge Electronic Design Ltd., Cambridge, UK) and CED Signal software (Version). The EMG was sampled at a rate of 1000 Hz with online 30 Hz high-pass, 1000 Hz low-pass, and 60 Hz notch filters. For GO-, STOP-, and IGNORE-signal trials, EMG sweeps were triggered 90 ms prior to each TMS pulse, and EMG was recorded for 1 s. In all participants, the active baseline pulse was delivered prior to the start of GO trials (i.e., during the initial fixation period). In most participants, the EMG sweep triggered 10 ms after commencement of the trial and the TMS pulse was delivered at 100 ms. However, an initial programming error omitted this 90ms delay on the baseline trials in the first 13 participants. Importantly, active baseline MEPs did not differ in amplitude between these 13 participants and the remaining 15 (see additional details in the subsequent paragraph). Thus, no distinction is made in the ensuing analyses.

MEP amplitude was extracted semi-automatically from the EMG trace using ezTMS, a freely available TMS preprocessing tool developed for use in MATLAB (Hynd et al., 2021). MEP amplitude was defined as the difference between the maximum and minimum amplitude 10 - 50 ms after the TMS pulse. In addition to the automatic determination of MEP amplitude, each trial was visually inspected for accuracy without knowledge of the specific trial type. Trials in which the MEP amplitude was less than .01 mV or the root mean square (RMS) of the EMG trace was greater than .01 mV were excluded from the MEP analyses. The first 80 ms of the EMG sweep were used to calculate the RMS power (i.e., for all GO, STOP, IGNORE, and most active baseline trials), whereas for the active baseline trials in the first 13 subjects, the baseline RMS-power was calculated for 81 to 160 ms after the TMS pulse. These RMS-EMG values (which were only used to identify baseline trials for exclusion for excessive noise) did not differ between the first 13 subjects and the remaining 15.

Mean amplitude MEPs were computed for each condition for each participant (as a function of Signal Type and Stimulation Time) as well as separately for successful and failed STOP trials at each Stimulation Time. These values were then normalized by dividing them by the mean active baseline MEPs. For each condition, the average number of trials were as follows: 206 (active baseline), 76 (GO/150), 75 (GO/175), 80 (GO/200), 51 (IGNORE/150), 52 (IGNORE/175), 46 (IGNORE/200), 28 (failed STOP/150), 28 (failed STOP/175), 26 (failed STOP/200), 27 (successful STOP/150), 26 (successful STOP/175), 26 (successful STOP/200). Data were excluded from analyses if, after combining successful and failed STOPs, any condition had fewer than 20 MEPs (which was the case for one participant). Additionally, for analyses that distinguished between successful and failed STOP trials, data were excluded if there were fewer than 10 MEPs in any condition (which was the case for two participants).

#### Data Analyses

The behavioral RT data were analyzed with RM-ANOVAs (with Huynh-Feldt correction in instances where the assumption of sphericity was violated) and Bayesian equivalents with non-informative priors in JASP software (Love et al., 2019). Planned and posthoc comparisons were made using paired t-tests with Holm-Bonferroni correction and Bayesian *t*-tests. Bayesian analyses were provided in addition to the frequentist statistics because some of our a priori hypotheses stipulated the presence of a null effect. The MEP data were analyzed principally by comparing each Signal Type at each Stimulation Time. Secondarily, we completed two-way ANOVAs in which Stimulation Time was considered as an additional factor, and these analyses are reported in Appendix 1.

Behavioral RT data for mean GO RT, mean IGNORE RT, and mean failed STOP RT were analyzed with a one-way RM-ANOVA. Errors were excluded from the analyses and were generally low for both GO and IGNORE trials (2 and 7%, respectively). Mean STOP signal reaction time (SSRT) was estimated according to the integration method with go-omission replacement, as detailed by Verbruggen et al. (2019). We also examined mean p(inhibit) and the difference in failed STOP and GO RT to ensure that the SST portion of our task functioned properly.

We completed a variety of analyses on the MEP data. First, and most critically for our prediction that MEP suppression would not differ for STOP and IGNORE signals at 150 ms, we compared mean MEPs for GO, STOP, and IGNORE signals at each Stimulation Time. Second, we likewise compared successful STOP, failed STOP, and IGNORE signals at each Stimulation Time. In both cases, we report the results of frequentist Paired t-tests (with Holm-Bonferroni alpha correction) alongside their Bayesian counterparts. Due to our null prediction, it should be noted that, when reporting Bayes Factor (*BF*), we report *BF*_01_ (evidence supporting the null hypothesis) or BF_10_ (evidence supporting the alternative hypothesis), depending on which way the evidence pointed.

Lastly, we complemented these contrast-based analyses with rm-ANOVA versions that included both Signal Type and Stimulation Time as factors. These results are contained in Appendix 1.

#### Open Data Practices

The data files, as well as scripts for running the task and analyzing the data can be found on the Open Science Framework (OSF) at https://osf.io/b72mp/?view_only=75676ea4492a41a59df79528e6f81b41.

## Results

### Behavior

The behavioral RT results for Experiment 1 are presented in Figure 2 *(left panel)*. The comparison for GO, IGNORE, and failed STOP RTs revealed a strong effect (*BF*_10_ = 4.73×10^33^, *F*(1.90, 51.27) = 148.72, *p* < .001, 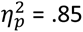). Posthoc comparisons indicated that participants responded faster to GO signals (mean: 541 ms, *SD =* 48.8) than to IGNORE signals (mean: 605 ms, *SD* = 66.6; *BF*_10_ = 2.30 x 10^9^, *t*(27) = 13.50, *p*_holm_ < .001, *d* = 2.55). Likewise, participants responded faster during failed STOP trials (mean: 529, *SD* = 57.2) than during IGNORE-signal trials (*BF*_10_ = 2.68 x 10^12^, *t*(27) = 16.05, *p* < .001, *d* = 3.03). Importantly, RT was also faster on failed STOP trials than on GO trials (*t* = 2.55, *p*_Holm_ = .014, *d* = .48, *BF*_10_ = 8.06), and the mean p(inhibit) was .495 (*SD* = .03). Thus, the assumptions of the race model held (e.g., Verbruggen et al., 2019), and the task was effective at maintaining a p(inhibit) around .5. The mean SSRT was estimated to be 305 ms (*SD* = 56.6).

**Figure 2.**
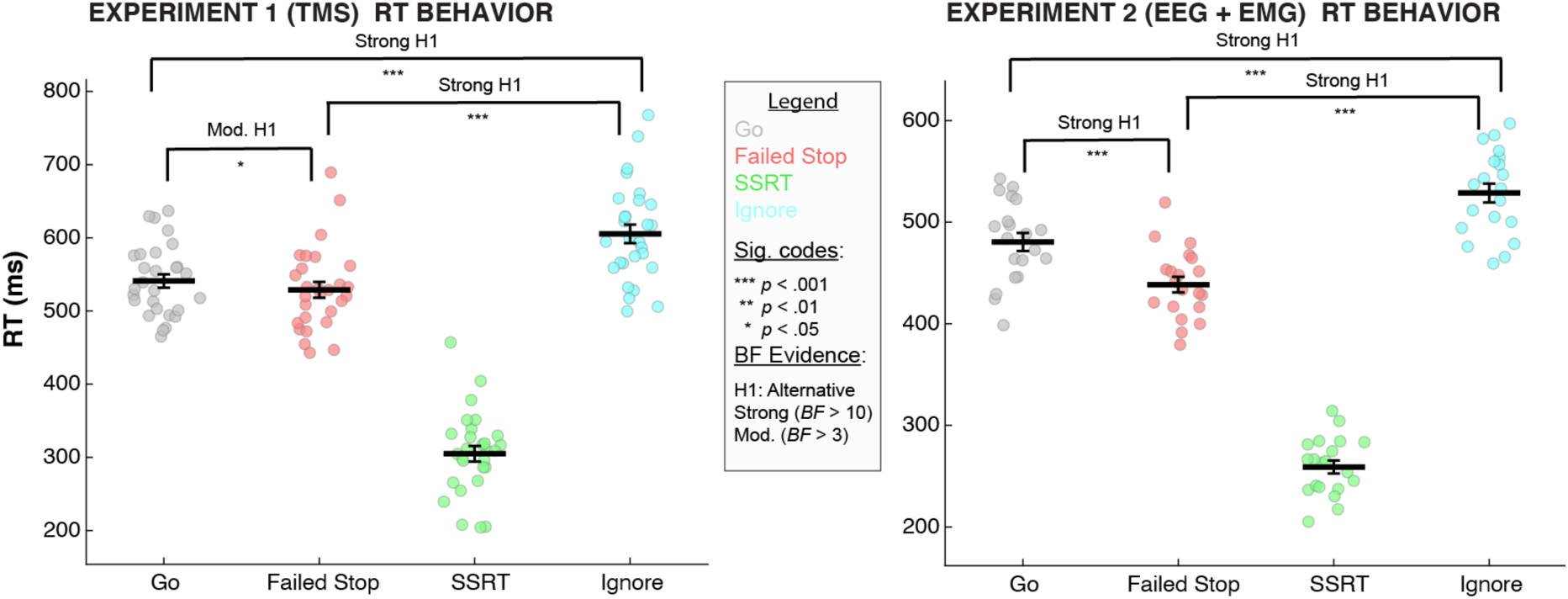
Behavioral RT results from Experiments 1 (left) and 2 (right). Horizontal black brackets indicate comparisons made using paired t-tests. We report both Frequentist (p-values with Holm-Bonferroni correction) and Bayesian (Bayes Factor) results. Horizontal black bars indicate the group mean RT whereas points represent individual participant mean RTs. Error bars denote SEM. SSRT was estimated via the integration method (Verbruggen et al., 2019) and not statistically compared to the other, directly observed RTs.

### Non-selective CSE suppression

We first aimed to replicate the established findings of non-selective CSE suppression in the task related muscle following both STOP (Badry et al., 2009) and IGNORE signals (Iacullo et al., 2020). As can be seen in Figure 3 (*left panel*), global MEP suppression was evident for both IGNORE and STOP signals at every Stimulation Time relative to GO signals (STOP/150: *BF*_10_ = 437.49, *t*(27) = 5.77, *p*_Holm_ < .001, *d* = 1.09; IGNORE/150: *BF*_10_ = 1098.09, *t*(27) = 5.29, *p*_Holm_ < .001, *d* = 1.00; STOP/175: *BF*_10_ = 8570.78, *t*(27) = 6.93, *p*_Holm_ < .001, *d* = 1.31; IGNORE/175: *BF*_10_ = 82.36, *t*(27) = 4.51, *p*_Holm_ < .001, *d* = .85; STOP/200: *BF*_10_ = 3315.45, *t*(27) = 5.42, *p*_Holm_ < .001, *d* = 1.02; IGNORE/200: *BF*_10_ = 10.60, *t*(27) = 3.56, *p*_Holm_ = .002, *d* = .67; also see Appendix 1 for the ANOVA main effect of Signal Type).

**Figure 3.**
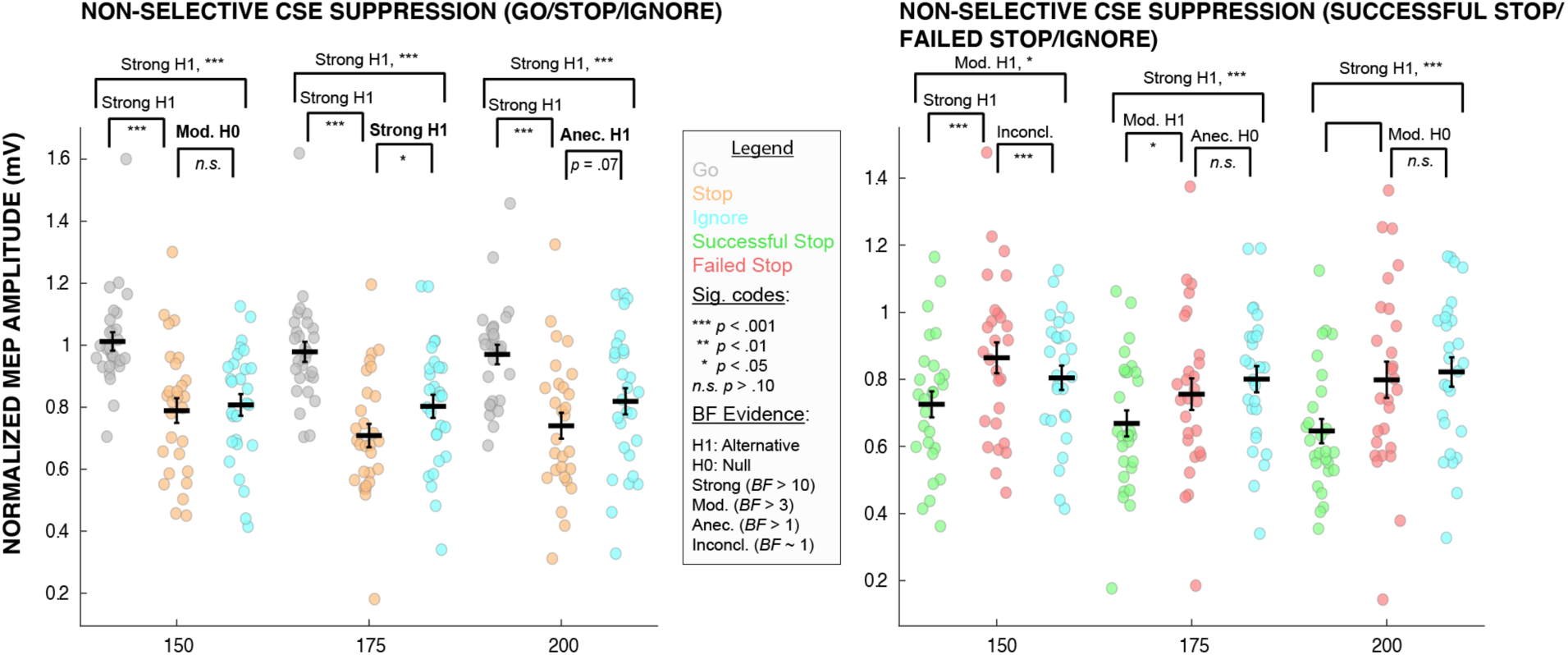
Mean normalized motor-evoked potentials for each Signal Type at each TMS Stimulation Time. Horizontal black brackets indicate comparisons made using paired t-tests. We report both Frequentist (p-values with Holm-Bonferroni correction) and Bayesian (Bayes Factor) results. Horizontal black bars indicate the group mean MEP whereas points represent individual participant’s mean normalized MEPs. Error bars denote SEM. (Left panel) Comparisons between Go, Stop (including both successful and failed stop trials), and Ignore trials. (Right panel) Comparisons between Successful Stop, Failed Stop, and Ignore trials.

More importantly for the present study, we then directly compared the MEPs for STOP and IGNORE signals to evaluate our prediction that the MEPs would be equivalent, at least at the earliest Stimulation Time of 150 ms. Critically, there was moderate evidence (*BF*_01_ = 4.00, *t*(27) = .48, *p*_Holm_ = .64, *d* = .09) that there was indeed no difference between STOP and IGNORE signals when CSE was measured 150 ms post-stimulus onset. By contrast, strong evidence for a difference between STOP and IGNORE signals started to emerge at 175 ms (BF_10_ = 15.40, *t*(27) = 2.42, *p*_Holm_ = .019, *d* = .46), with STOP signals exhibiting greater global MEP suppression. Although there was also numerically greater MEP suppression for STOP signals at the 200 ms Stimulation Time, there was only anecdotal evidence to support this difference (*BF*_10_ = 1.28, *t*(27) = 1.86, *p*_Holm_ = .068, *d* = .35). Importantly, these results confirm our hypothesis that while both STOP and IGNORE signals produce non-selective CSE suppression, this effect does not initially differ until later in the trial.

When STOP trials were distinguished by stopping success, further interesting patterns emerged. First, when distinguishing between successful and failed STOP trials (see Figure 3, *right panel*), global MEPs were clearly reduced for successful STOP trials at all Stimulation Times, again replicating prior work (Badry et al., 2009; Wessel et al., 2013; Jana et al., 2020). Strong evidence for this difference was present at 150 ms (*BF*_10_ = 173.88, *t*(26) = 4.61, *p*_Holm_ < .001, *d* = .89) whereas the evidence was Moderate at 175 and 200 ms (respectively: *BF*_10_ = 2.10, *t*(26) = 2.51, *p*_Holm_ = .030, *d* = .48; *BF*_10_ = 6.28, *t*(26) = 3.43, *p*_Holm_ = .002, *d* = .66).

We also found greater MEP suppression for successful STOP relative to IGNORE trials at all stimulation times, with Moderate evidence at 150 ms (*BF*_10_ = 3.91, *t*(26) = 2.63, *p*_Holm_ = .023, *d* = .51) and Strong evidence at 175 and 200 ms (respectively: *BF*_10_ = 360.25, *t*(26) = 3.81, *p*_Holm_ = .001, *d* = .73; *BF*_10_ = 212.95, *t*(26) = 3.95, *p*_Holm_ < .001, *d* = .76). For failed STOP relative to IGNORE trials, it was inconclusive whether there was evidence for a difference or not at 150 ms (*BF*_10_ = 1.02, *t*(27) = 1.99, *p_Holm_* = .052, *d* = .38), although it is notable that MEPs were numerically higher for failed STOP trials. However, there was Anecdotal evidence for no difference between failed STOP and the IGNORE trials at 175 ms (*BF*_01_ = 2.62, *t*(26) = 1.30, *p*_Holm_ = .200, *d* = .25) and Moderate evidence for no difference at 200 ms (*BF*_01_ = 4.26, *t*(26) = .52, *p*_Holm_ = .606, *d* = .10).

In summary, these results suggest that IGNORE and STOP trials feature equivalent amounts of non-selective CSE suppression at the early post-signal latency (here, 150ms), with successful STOP trials representing the part of the STOP trial CSE distribution that happened to contain more suppression compared to failed STOP trials. This suggests that this early activity contributes to the success of stopping, but is not unique to STOP trials. Moreover, these results suggest that in case of a STOP signal, this initial suppression is sustained further into the post-signal period, as significant differences to IGNORE trials emerged at later latencies (here, starting at 175ms).

## Experiment 2

### Method

#### Hypothesis

The hypotheses for this follow-up experiment were registered after the results of Experiment 1 were known. The pre-registration document (as well as data, task code, and preprocessing/analysis scripts) can be found at https://osf.io/9uqwa/?view_only=fa3603365ecb4a84b46e285a5436b986

Based on the findings in Experiment 1, we made two predictions:

1. The suppression of the overt EMG that is observable on many successful STOP trials (Raud & Huster, 2017; Raud et al., 2020; Jana et al., 2020) is not unique to STOP trials, but will instead also occur – to the same degree – on IGNORE trials.
2. Multi-variate pattern analysis (MVPA) of whole-scalp EEG will not show any significant differences between STOP and IGNORE signals in the early post-signal time period. However, in line with the CSE results from Experiment 1, we predicted that STOP and IGNORE signals would become decodable from one another at around 175ms.

#### Participants

Twenty healthy, right-handed adults (16 female, mean age: 22.6, *SD* = 2.7) participated in the EEG experiment after providing written informed consent. The participants were recruited via an email sent out to the University of Iowa community. Participants were compensated at an hourly rate of $15 or received course credit. The study was approved by the University of Iowa’s Institutional Review Board (#201511709).

#### Task

The task was identical to Experiment 1, with the following exceptions: 1) Responses were made using both index fingers (‘z’ for left and ‘m’ for right on a QWERTY keyboard). 2) The total number of trials was reduced to 720 trials that were separated into ten blocks (66.7% GO, 16.7% STOP, and 16.7% IGNORE trials).

#### EEG and EMG recording

EEG was acquired using a Brain Product ActiChamp 63-channel system. The ground and reference electrodes were placed at AFz and Pz, respectively. EMG was recorded from each hand using two electrodes placed on the belly and tendon of each FDI muscle. Ground electrodes were placed on the distal end of each hand’s ulna. The EMG electrodes were connected to the EEG system using two auxiliary channels (via BIP2AUX adaptor cables). Thus, there were 65 total recording channels. All channels were sampled at a rate of 2500 Hz. To minimize task-unrelated EMG activity, the participants were instructed to only move their index fingers when necessary and to otherwise keep both hands relaxed and pronated on the desk.

#### EEG and EEG preprocessing

Custom MATLAB code was used to preprocess the EEG and EMG data. The EMG data analysis was adapted from Raud & Huster (2017). Both EEG and EMG data were bandpass filtered (EEG: 0.5-50Hz; EMG: 2-200Hz) and downsampled to a rate of 500Hz. The continuous EEG data were visually inspected to identify non-stereotypical artifacts. Segments with non-stereotypical EEG artifacts were removed from both the EEG and EMG data. The EEG data were re-referenced to the common average and submitted to an infomax independent component algorithm (Makeig et al., 1996) as implemented in the EEGLAB toolbox (Delorme & Makeig, 2004). The resulting independent components were visually inspected to remove components that captured stereotypical artifacts (namely, blinks and saccades). Both the EEG and EMG data were epoched relative to the onsets of GO, STOP, and IGNORE stimuli (-200 – 1000ms). For GO and IGNORE trials, all trials with incorrect or absent responses were excluded from all analyses. The EMG data were then converted to root mean square power using a sliding window of +/- 5 sample points and baseline corrected by dividing the entire epoch by the mean of the 200ms pre-stimulus EMG. The resulting data were then standardized for each hand across all types of trials and sample points using the z-transform.

#### Behavior Analysis

Behavior data were analyzed identically to the behavior data in Experiment 1.

#### EMG validation

As a manipulation check, we examined whether the EMG data accurately reflected motor output by correlating the latency of each trial’s GO trial EMG peak to GO RT on the same trial within each subject. If EMG and GO RT are meaningfully related as predicted, the mean individual-subject correlation should be significantly larger than zero. Pearson correlation coefficients (*r*) were calculated for each subject at the single-trial level. For those correlations, trials in which Cook’s *D* exceeded 4/N were deemed statistical outliers and were not included in calculating the individual *r*’s (on average, 6.2% of trials per subject). To ensure normality, the individual *r*’s were transformed using Fisher’s z-transformation. The individual, z-transformed *r* values were then submitted to a one-sample *t*-test against 0 on the group level. To enable their original interpretation as correlation coefficients, the averaged z-transformed *r* values were then transformed back to *r* (see Silver & Dunlap, 1987).

#### Partial EMG (prEMG) trial identification

Previous research has shown that residual EMG on successful STOP trials shows a marked reduction around 150 ms following the STOP signal (Raud & Huster, 2017; Raud et al., 2020). We here hypothesized that the same would be true for IGNORE trials. This analysis was again adapted from Raud & Huster (2017). For each subject, we limited the time window of successful STOP EMG traces to 0-200ms and averaged the time-constrained EMG traces across trials. Using the subject-specific EMG traces, a detection threshold was identified by finding the peak amplitude of the averaged EMG trace in the first 200ms following the signal for each subject (mean threshold (*z*-score) = 0.3, *SD* = 0.39). Within each subject, a trial was then denoted a prEMG trial if the individual-trial peak in the same window exceeded the threshold. The same procedure was used to identify prEMG on IGNORE trials, except that all IGNORE trials with correct responses were used, as they all contained prEMG activity.

#### EMG analysis

Figure 4 (*upper left*) depicts the grand-averaged EMG traces for successful STOP, failed STOP, and IGNORE trials, separately for the selected and unselected hands. Statistical analyses were performed on EMG data from the selected hand. Both successful STOP and IGNORE trials showed an initial pattern of ramping activity, which was, around 150ms following either signal, interrupted by a sudden downturn. In STOP trials, the EMG trace subsequently returned towards zero, in line with the outright cancellation of the response. In IGNORE trials, the downturn was followed by a second ramping, in line with a reactivation of the response. Consistent with prior studies using only STOP trials, the variable of interest was the latency of the downturn after the initial ramping, which is taken as an indicator of early inhibitory activity (Raud & Huster, 2017; Raud et al., 2020; Jana et al., 2020). To this end, the peak of the EMG trace was detected within the first 200ms after successful STOP and IGNORE signals and its latency averaged for each subject and condition separately. We refer to this latency, which was the DV of interest for the EMG analysis, as the prEMG-int (partial EMG interruption) latency.

**Figure 4.**
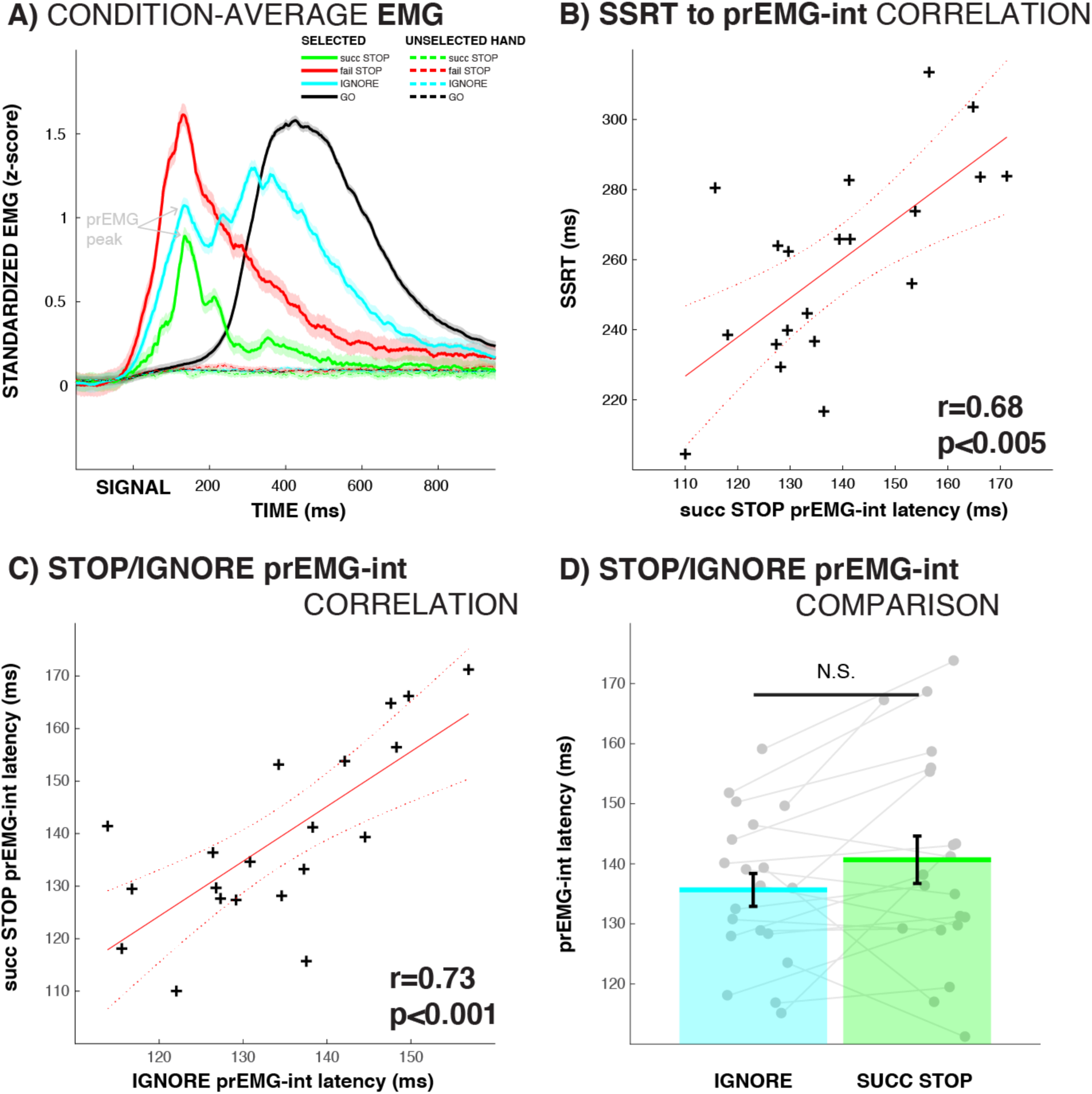
EMG data results. Upper left (A): EMG traces for all conditions. Arrows on the left portion of the plot indicate prEMG-int latency for the successful STOP and IGNORE condition. Upper right (B): group level correlation between successful STOP prEMG-int latency and SSRT. Lower left (C): group level correlation between successful STOP and IGNORE prEMG-int latency. Lower right (D): successful STOP and IGNORE prEMG-int latency comparison.

To test hypothesis #1 for Experiment 2 (see Hypothesis section above), first, we correlated mean STOP trial prEMG-int latency with IGNORE trial prEMG-int latency at the group-level to examine the extent to which both measures related to each other. Furthermore, we compared both latencies with a paired *t*-test. Since we hypothesized that the same interruption that has been observed on STOP trials would also occur on IGNORE trials, we predicted that these latencies would be highly correlated across subjects and that their condition means would not differ. As with Experiment 1, we report *BF* alongside frequentist statistics for all EMG analyses in Experiment 2.

We also examined the relationships between prEMG and behavioral RT measures. First, we correlated STOP prEMG measures with mean SSRT to replicate prior work that showed a strong correlation between the two (e.g., Raud & Huster, 2017). For IGNORE trials, we used the same single-trial level approach to correlate prEMG measures with RT that was described for the GO trial analysis in the EMG validation section above. To provide an unbiased estimator of IGNORE RT, it was quantified from the onset of the IGNORE signal, not of the GO signal (though the results were qualitatively unchanged when the full RT was taken instead).

#### EEG Multi-variate pattern analysis

To test hypothesis #2 for Experiment 2, we investigated at which time point a multi-variate pattern analysis (MVPA) classifier applied to the whole-scalp EEG data recorded during the task would be able to distinguish between the conditions of interest above chance, with a particular focus on the first time point at which whole-scalp EEG showed above-chance decoding of STOP vs. IGNORE trials.

As control analyses (and to illustrate the performance of the classifier approach), we also examined time points during which successful STOP, failed STOP, and IGNORE trials differed from matched GO trials. To do so, GO trials were split into fast and slow GO trials by each subject’s median GO RT for each hand. Fast GO trials were matched to failed STOP trials (which represent the fast part of the GO RT distribution, Logan et al., 1984) and slow GO trials were matched to successful STOP and IGNORE trials (since successful STOP trials represent the slower part of the RT distribution, and IGNORE RT was longer than regular GO RT; cf. behavioral results for Experiments 1 and 2). This matching procedure ensured that decoding was not confounded by differences in RT. In addition, the number of left and right arrow trials were balanced for all decoding analyses to ensure non-biased results.

MVPA analyses were performed using the ADAM toolbox (Fahrenfort et al., 2018). We used leave-one-out cross-validation in which a Linear Discriminant Analysis classifier was trained on all trials but one and tested on the remaining trial that was not part of the training set. This validation was implemented in the sample-point-wise fashion, in which training and testing were done at each sample point following the GO/STOP/IGNORE signal. Classifier performance was measured as the area under the receiver operating characteristics curve (AUC; Wickens, 2002), which quantified the total area of the curve when the cumulative true positive rate (probability of correct classification) was plotted against the false positive rate (probability of incorrect classification) for each decoding problem between the conditions of interest.

Time series of AUC values were obtained for each pair of conditions and each participant. To identify time points during which classification performance was significantly above chance, we compared each AUC value against chance-level performance (0.5) using one-sample *t*-tests, which was corrected for multiple comparisons using cluster-based permutation testing (10000 iterations, cluster-p value=.00001, individual sample point alpha=.0001; Maris & Ostenveeld, 2007).

#### Event-related potential analysis

Since the fronto-central ERPs are commonly associated with stop-signal performance (e.g., de Jong et al., 1990; Huster et al., 2020), we also quantified the condition ERPs at the fronto-central electrodes Cz and FCz. All epochs were baseline corrected by subtracting the mean of the 200ms pre-stimulus period. Differences between the conditions of interest (successful STOP vs. IGNORE, failed STOP vs. IGNORE, and successful STOP vs. failed STOP) were tested for significance using sample-point-wise paired samples *t*-tests, in which the resulting *p*-values were corrected for multiple comparisons using the false-discovery rate (FDR) correction procedure (Benjamini & Hochberg, 1995).

## Results

### Behavior

The RT results for the EEG experiment, which involved hand responses, paralleled those of Experiment 1, which was performed using the feet (with slightly faster RT values across the board, see Figure 2, *right panel*). There was again a strong effect of RT, *BF*_10_ = 6.84 x 10^14^, *F*(1.73, 32.80) = 134.25, *p* < .001, 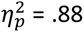. As with Experiment 1, RT was faster to the GO signal (mean = 480 ms, *SD* = 39.8) than to the IGNORE signal (mean 529 ms, *SD* = 41.7; *BF*_10_ = 5.35 x 10^5^, *t*(19) = 8.75, *p*_Holm_ < .001, *d* = 1.96. RT was also faster to failed STOP trials (mean = 438, *SD* = 34.4 ms) than to IGNORE trials (*BF*_10_ =1.33 x 10^9^, *t*(19) = 16.37, *p*_Holm_ < .001, *d* = 3.66) and faster to failed STOP than to the GO trials (*BF*_10_ = 6.57 x 10^6^, *t*(19) = 7.62, *p*_Holm_ < .001, *d* = 1.70). The mean p(inhibit) was .5 (*SD* = .016), and the mean SSRT was 259 ms (*SD* = 28.3). Thus, the task was effective at producing approximately equal numbers of successful and failed stops, and the assumptions of the independent race model held (Verbruggen et al., 2019).

### EMG Validation

Single-trial correlation analysis revealed a strong positive correlation between GO trial EMG peak latency and GO RT (mean Pearson’s *r =* 0.85, *BF*_10_ = 1.30×10^7^, *t*(19) = 11.21, *p* < 10^-9^, *d* = 2.51), validating the relationship between single-trial EMG and behavior.

### EMG

Grand-averaged EMG traces showed that there was an early EMG peak (around 140 ms) following both the STOP and IGNORE signals, which was followed by a sudden downturn (Figure 4, *upper left panel*). In line with our expectations, after this initial dip, the IGNORE EMG trace promptly began rebounding (around 200 ms), whereas the successful STOP EMG trace became further suppressed.

Further in line with our hypothesis that both STOP and IGNORE trials would show a comparable early-latency suppression of EMG (Hypothesis #1), successful STOP and IGNORE trial prEMG-int latencies showed a strong positive correlation (Pearson’s *r* = .73, *BF*_10_ = 141.26, *p* < .001) and were not statistically different from one another, though evidence for this was inconclusive (*t*(19)=-1.88, *p* = .076, *d* = 0.42, *BF*_01_ =1.01,). If anything, prEMG-int latency was numerically longer on successful STOP trials compared to IGNORE trials.

In line with prior studies, successful STOP trial prEMG-int latency (mean latency = 139 ms, *SD* = 17.2) was significantly positively correlated with SSRT (*r* = .68, *BF*_10_ = 40.79, *p* < .005). Moreover, both IGNORE trial prEMG-int latency (mean Pearson’s *r* = .22, *BF*_10_ = 64.84, *t*(19) = 4.17, *p* < .001, *d* = 0.93) and its peak amplitude (mean Pearson’s *r* = -0.46, *BF*_10_ = 2.96×10^6^, *t*(19) = - 10.18, p < .001, *d* = 2.28) were significantly correlated to IGNORE RT.

In summary, the EMG data from Experiment 2 show that both IGNORE signals and STOP signals are followed by low-latency interruptions of the EMG. While this pattern on STOP trials is typically interpreted as a unique signature of outright action-stopping, our analyses indicate that this is a universal signature of inhibition that is common to all salient events. Successful STOP trials did not incur faster EMG suppression compared to IGNORE trials, and the latencies of the prEMG-int latency was highly correlated across subjects, suggesting that the same process is active on both types of trials.

### MVPA

The results of the MVPA analysis are presented in Figure 5, alongside the fronto-central ERPs. In line with our hypotheses and our findings from Experiment 1, IGNORE trials and successful STOP trials could not be successfully decoded from one another until 180 ms after signal onset (periods of above-chance decoding: 180-792 ms, 820-880 ms). IGNORE trials and failed STOP trials could not be decoded from one another until 428 ms following signal onset (significant period: 428 - 758 ms).

**Figure 5.**
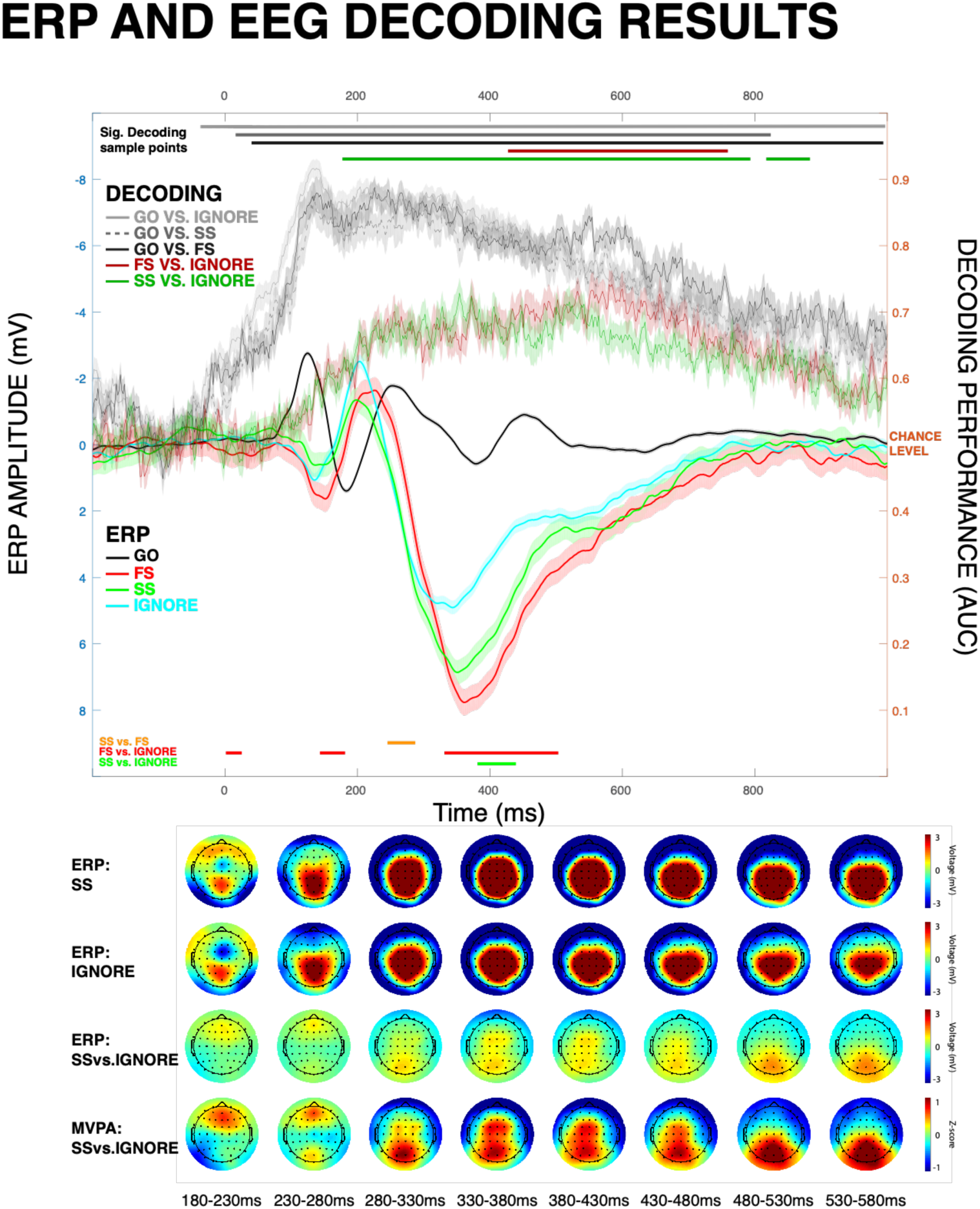
ERP and EEG DECODING results. Upper plot contains the condition ERPs at fronto-central electrode sites FCz/Cz (left scale) and decoding performance for the contrasts of interest (right scale, 0.5 = chance-level decoding). Color bars above the figure denote time periods with significant above-chance whole-scalp MVPA decoding (color-coded identically to the legends). Color bars below the figure denote significant differences in the fronto-central ERP (orange: SS vs. FS; red: FS vs. IGNORE; and green: SS vs. IGNORE). The lower plot shows topographical distribution for SS, IGNORE, and SS-IGNORE ERP; and forward-estimated decoding estimates of SS vs. IGNORE decoding.

Notably, when the classifier was trained to distinguish between GO and other trials (failed STOP, successful STOP, and IGNORE), decoding performance was significantly above chance for almost the entirety of the epoch (see Figure 5). This suggests that our analysis methods were not too conservative to detect early differences in the contrasts of interest. One interesting anecdotal observation is that classifier performance peaked around 140 ms for all combinations of STOP/IGNORE vs. GO comparisons. This coincides with the time period during which all three trial types (successful/failed STOP, IGNORE) showed CSE suppression for STOP and IGNORE trials in Experiment 1.

### ERP

Both successful and failed STOP trials showed significantly stronger positive voltage deflection compared to IGNORE trials during latencies corresponding to the P3 (successful: 388-436 ms; failed: 336-506 ms). No significant differences were found in the N2 time range (see Figure 5). Indeed, if anything, the N2 amplitude was larger for IGNORE trials compared to both STOP trial types. There were earlier ERP differences between failed STOP trials and IGNORE trials (2-22 ms and 148-182 ms), but not for successful STOP and IGNORE trials. During both periods, failed STOP trials showed increased positivity relative to IGNORE trials, perhaps indexing motor processes. Finally, the successful vs. failed STOP ERP comparison showed significant differences around SSRT (250-288 ms; SSRT = 259 ms) reflecting the fact that, in line with prior literature, successful STOP trials showed an earlier onset of the P3 compared to failed STOP trials (Kok et al., 2004; Wessel & Aron, 2015).

These findings show that late-latency fronto-central ERPs (viz., the fronto-central P3) uniquely reflect processes related to outright stopping. In comparison, the fronto-central N2 was, if anything, larger for IGNORE trials. No earlier differences between successful STOP and IGNORE trials were found.

## Discussion

In this study, we tackled what are arguably the two most prominent debates in the recent literature on action-stopping: the question of which processes reflect the attentional detection of a STOP signal and which index the actual implementation of inhibitory control, as well as the question of why purported inhibitory signatures occur at two different latencies following a STOP signal. Our results support the following view: Early signs of motor inhibition following STOP signals – including EMG suppression and early non-selective CSE reduction – are not unique to action-stopping, and indeed occur after any type of salient event, notably to the same degree. Differences between outright movement cancellation (on STOP trials) and attentional capture (on IGNORE trials) only emerge after this initial inhibitory response. Specifically, the EEG-MVPA analysis in Experiment 2 revealed that STOP- and IGNORE-trials are only reliably decodable from one another from around 180 ms onwards. This is in line with the divergence in CSE suppression at the level of corticomotor tracts, which emerge at 175 ms. There is no evidence for differences in prior signatures (CSE, EMG, EEG). Notably, the early-latency EMG suppression that has been proposed as the earliest sign of inhibition in several recent studies (Raud & Huster, 2017; Huster et al., 2020; Jana et al., 2020) is not unique to outright action-stopping and occurs at the same latency and with the same magnitude after IGNORE-signals as well.

There are several notable features of these data, which inform the basic research on action-stopping across methodologies. First, with regards to scalp EEG, our results show that the earliest signals that distinguish stop- from ignore trials occur at around 180 ms following the STOP signal. Moreover, the ERP at the fronto-central electrodes (which provide the strongest predictive power in the forward weight projection of the MVPA analysis during that time window) indicates that this initial decoding difference is actually due to an increase of the fronto-central N2 ERP in the IGNORE relative to the STOP condition. The first signature at which both decoding weights and ERP difference indicate a unique increase of activity on successful STOP vs. IGNORE trials is the fronto-central P3. This is in line with the proposal that this ERP reflects a process that is unique to outright action-stopping (de Jong et al., 1990; Kok et al., 2004), and with the proposal that the onset latency of this ERP index the onset of processing that distinguishes successful from failed stopping (Wessel & Aron, 2015). Second, with regards to CSE suppression, these results confirm earlier studies showing that such effects can be observed not just after STOP signals, but also after surprising or merely infrequent events (Wessel & Aron, 2013; Dutra et al., 2018; Iacullo et al., 2020). The current results substantially add to these findings by demonstrating that STOP signals and IGNORE signals do not differ in the degree to which they suppress CSE early on (here, at 150ms post-stimulus). However, there is divergence in this suppression at later time points, in that STOP signals do begin to show additional suppression at 175ms. This suggests that in the case of STOP signals, additional processing sustains the initial automatic CSE suppression caused by the saliency of the STOP signal. It is tempting to assume that this additional suppression is due to the process that is indexed by the EEG signatures that emerge in the MVPA analysis during roughly the same time period. Moreover, another interesting pattern in the CSE suppression results is that despite the absence of differences in overall stop- and ignore-trial amplitudes at the early 150ms post-stimulus time point, there is already a significant difference between successful and failed stop-trials. This suggests that while early CSE suppression is due to processes related to the attentional detection of the STOP signal, those processes do contribute to the success of outright stopping. Indeed, it appears that successful stop-trials at least partially reflect merely the stochastic subsample of trials in which the initial attention-related CSE suppression just happens to be stronger. Hence, successful action-stopping likely results from a combination of early-latency inhibitory processes triggered by the attentional detection of the STOP signal and slower processes that are unique to STOP trials. This two-stage cascade is in line with a recent proposal by Schmidt & Berke (2017, see next paragraph). Lastly, with respect to EMG suppression, the current results are the first to demonstrate that this signature is not uniquely indicative of inhibitory control in the context of outright action-stopping, but is instead likely common to all salient events, further in line with the assertion that the automatic invocation of inhibitory control is a ubiquitous consequence of stimulus-driven attentional capture (Wessel & Aron, 2017). As briefly mentioned above, in the wider context of the cognitive neuroscience of motor inhibition and action-stopping, the current results support the view that action-stopping is a two-stage process. The first stage contains the automatic invocation of inhibitory control that is common to all salient events. This stage is followed by the – perhaps controlled – invocation of additional processes that are unique to action-stopping. The timing of processes in both the CSE suppression data of Experiment 1 and the EEG-MVPA data of Experiment 2 suggest that the latter processes do not emerge until around the 175 ms mark after the STOP signal (at which point both CSE and EEG-MVPA indicate differences between stop- and ignore signals). In our data, the first stage is evident as early as ∼140ms following signal onset, which was the average latency of the onset of EMG suppression. Based on these results and recent work in both human and non-human animals, we propose that the neural cascade underlying human action-stopping roughly corresponds to recently proposed models that postulate a two-stage stopping process (such as the pause-then-cancel model of Schmidt & Berke, 2017; Schmidt et al., 2013). Primarily based on rodent work on basal ganglia circuitry, their model proposes that outright action-stopping is achieved by a combination of two processes: an early-latency “pause” process that leads to an initial inhibition of the go-process, followed by a slower “cancel” process, which removes the ongoing energization of the go-response. The two processes operate at different latencies, but interact insofar as the implementation of the “pause” process buys time for the cancel process to achieve successful action-stopping (see also Wiecki & Frank, 2013 for a similar proposal of a broad, non-selective “pause” response at early, post-signal latencies). Adapting this framework to the human domain, we propose that the early signatures that are common to both STOP- and IGNORE signals – viz., early CSE / EMG suppression and EEG activity prior to 180ms – reflect the “pause” stage of this model. Indeed, the observed suppressive effects on EMG and CSE are in line with a suppression of the active go-process after salient events. We further propose that later activity – EEG activity in the time range of the fronto-central P3 and sustained CSE suppression beyond the early-stage – reflect the cancel stage. Indeed, such an adaptation of the pause-then-cancel model is in line with a large body of existing work in humans that goes beyond the current study. Specifically, we propose that the initial “pause” phase is implemented via the rIFC-STN hyper-direct pathway that has recently been empirically described in humans (Chen et al., 2020) and has long been proposed to be underlying a substantial part of the inhibitory activity involved in action-stopping (Aron et al., 2007). However, we suggest that the “pause” phase is not specific to action-stopping, but indeed ubiquitous to any type of salient event that captures attention (Wessel & Aron, 2017). This perspective unites the ostensibly disparate notions of rIFC as a “circuit breaker” (Corbetta & Shulman, 2002) or non-inhibitory detector of infrequency (Erika-Florence et al., 2014) and rIFC as an inhibitory node (Aron et al., 2007; 2014). As such, we propose that rIFC instantiates an inhibitory “pause” response, but not one that is unique to action-stopping. Physiologically, this “pause” response results in, amongst other things, early non-selective suppression of CSE, as well as suppression of any overt electromyographic activity. While the strength of this incidental invocation of inhibition following attentional detection can contribute to the success of action-stopping, it is not unique to outright action-stopping. We then further propose that in situations that do require outright action-stopping, a second “cancel” phase is implemented by brain structures underlying the fronto-central ERPs, which likely includes the pre-SMA (Swann et al., 2011; Enriquez-Geppert et al., 2010). We propose that this second phase consists of a broad retuning of active motor plans that are geared towards the exact behavioral requirements posed by the specific signal that initially triggered the pause-then-cancel cascade. In the case of outright action-stopping after STOP signals, this would consist of a shutting down of any still-active drive towards the execution of the motor response (Schmidt & Berke, 2017). In situations in which the salient signal does not instruct outright action-stopping, this second phase could still consist of more subtle adaptations of motor output. This would explain, amongst other things, why fronto-central EEG activity in the P3 time range relates to – for example – elongations in reaction time following unexpected (Wessel & Huber, 2019) or infrequent events (Waller et al., 2019), as well as changes in isometric force following such events (Novembre et al., 2018; 2019). As such, we propose a broader adaptation of the pause-then-cancel model of Schmidt and Berke (which is specific to the stop-signal task): a pause-then-*refine* mechanism that is shared by all infrequent events (with *refine* corresponding to *cancel* in the specific case of the stop-signal task). The full theoretical implications of such a model in the context of the human brain still have to be explicitly formulated, but we do believe that such a model is favored by the majority of available research, not least the current study.

One potential point of criticism of the current work could be the notion that since the infrequent IGNORE events in the current experiment were presented within the context of the stop-signal task, participants may have been biased towards using inhibitory control after their occurrence. Indeed, some behavioral work indicates that participants may employ a “stop-then-discriminate” strategy when facing infrequent events while also anticipating stop-signals (Bissett & Logan, 2014). This work is not at odds with our interpretations, and we in fact believe the “stop-then-discriminate” strategy to be another term for the ubiquitous “pause” phase in Schmidt & Berke’s, or the “Hold your horses” conceptualization of Frank and colleagues (Frank, 2006 ; Frank et al., 2007). Moreover, there is clear convergent evidence that IGNORE events, even when presented outside of a stop-signal task context (and hence without proactive control), produce non-selective CSE suppression (Iacullo et al., 2020), fronto-central EEG activity (Courchesne et al., 1975; Wessel & Huber, 2019; Novembre et al., 2018; 2019), slowing of motor behavior (Waller et al., 2019), changes in isometric motor activity (Novembre et al., 2018; 2019), and rIFC activity (Corbetta & Shulman, 2002; Erika-Florence et al., 2014). Hence, the observed CSE, EMG, and EEG changes here are not contingent on the presence of proactive control. Instead, the current study shows that when proactive control requirements are matched, both STOP and IGNORE signals produce the same degree of early-latency inhibitory activity. A two-task solution in which participants would have first performed a task that included IGNORE events but no STOP signals (followed by a separate stop-signal task session), would not have allowed for any clear insights using the EEG-MVPA approach used here, precisely because the presence of proactive control in only the stop-signal task would have produced broad changes to the EEG activity (Kenemans, 2015; Elchlepp et al., 2016; Soh et al., 2021), which would have rendered the attempted MVPA approach in the current study moot. In sum, we believe that the current task produces an ideal comparison between IGNORE and STOP trials under conditions of matching degrees of proactive control. Since other research has clearly shown that the early-latency markers of inhibitory control that are common to STOP and IGNORE signals are also present when IGNORE signals occur in the absence of proactive control (see Wessel, 2018 for a review), we believe that our results cannot be explained by the presence of proactive control alone.

In summary, the current study used a comparison between the brain processes triggered by STOP signals and IGNORE signals. It enabled us to show that there are no meaningful differences in neural or physiological processing between both signals in any of the three domains (EMG, CSE, EEG) until around 175/180ms following STOP signals. Moreover, we have shown that IGNORE signals not only produce CSE suppression on par with STOP signals, but that the suppression of EMG that has recently been proposed as a trial-to-trial marker of the speed of the overall stopping process in the STOP signal task also occurs after merely infrequent events. As such, we believe that these data warrant a reexamination and reinterpretation of existing work in the domain of motor inhibition along the lines of recently proposed two-stage models of action-stopping – with the additional prediction that only the second stage of such models is actually unique to situations that require outright action-stopping.

## Acknowledgements

This work was funded by grants from the National Science Foundation (NSF CAREER 1752355 to JRW) and the National Institutes of Health (NIH R01 NS102201 to JRW).

## Conflict of interest

The authors report no financial conflict of interest.

## Appendix 1

Although we primarily examined CSE results by comparing the mean MEPs for different trial types separately at each time point (e.g., paralleling how we analyzed EEG data), we additionally examined CSE by submitting the mean normalized MEPs to RM-ANOVAs with the within-subject factors of Signal (GO, STOP, and IGNORE) and Stimulation (150, 175, 200). There was a main effect of Signal Type, *F*(1.58, 42.68) = 32.61, *p* < .001, 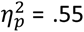, *BF*_10_ = 4.76 x 10^20^. There was also a main effect of Stimulation Time, *F*(1.96, 52.93) = 3.65, *p* = .033, 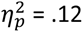, *BF*_01_ = 8.85, although the Bayesian analysis provided moderate evidence against the null hypothesis that Stimulation Time differed. There was no significant Signal Type X Stimulation Time interaction, *F*(2.97, 80.08) = 1.02, *p* = .39, 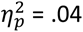, *BF*_01_ = 14.71. Post-hoc tests for Signal Type showed that MEPs were reduced for both IGNORE and STOP signals relative to GO signals [IGNORE: *t*(27) = 5.73, *p* < .001, *d* = 1.08, *BF*_10_ = 2.39 x 10^7^; STOP: *t*(27) = 7.79, *p* < .001, *d* = 1.47, *BF*_10_ = 1.60 x 10^12^].

As a whole, STOP signal MEPs were reduced compared to IGNORE signals, *t*(27) = 2.07, *p*_Holm_ = .044, *d* = .39, BF_10_ = 30.73, although it is worth reiterating that this difference appears to be driven primarily by the observed difference between STOP and IGNORE MEPs at 175 ms (and possibly 200 ms), whereas consistent with our hypothesis, no differences between IGNORE and STOP MEP suppression were found at the 150 ms Stimulation Time. Post-hoc tests for Stimulation Time indicated that 175 ms showed greater suppression than at 150 ms, *t*(27) = 2.65, *p*_Holm_ = .031, *d* = .50, *BF*_10_ = 2.40. There was no difference in MEPs at 150 vs. 200 ms, *t*(27) = 1.78, *p*_Holm_ =.162, *d =* .37, *BF*_01_ = 3.14, and no difference in MEPs at 175 and 200 ms, *t*(27) = 1.07, *p*_Holm_ = .29, *d* = .20, BF_01_ = 6.21.

We also completed the above analyses further distinguishing between successful and failed STOP trials. There was a main effect of Signal, *F*(2.25, 58.50) = 31.04, *p* < .001, 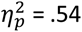, *BF*_10_ = 1.29 x 10^26^, and a main effect of Stimulation Time, *F*(1.67, 43.48) = 4.71, *p* = .013, 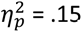, *BF*_01_ = 5.49 (again, Bayesian analyses provided moderate evidence against this main effect). There was no significant interaction, *F*(4.20, 109.10) = 1.68, *p* = .157, 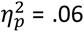, *BF*_01_ = 12.77. Posthoc tests for Signal Type indicated that successful STOP, failed STOP, and IGNORE MEPs were all reduced relative to GO MEPs [successful STOP: *t*(26) = 9.59, *p*_Holm_ < .001 , *d* = 1.85, *BF*_10_ = 3.11 x 10^19^; failed STOP: *t*(26) = 5.56, *p*_Holm_ < .001, *d* = 1.07, *BF*_10_ = 1.20 x 10^5^; IGNORE: *t*(26) = 5.47, *p*_Holm_ < .001 , *d* = 1.05, *BF*_10_ = 8.48×10^6^]. In addition, successful STOP MEPs were further reduced compared to IGNORE and failed STOP MEPs [respectively: *t*(26) = 4.13, *p*_Holm_ < .001, *d* = .79, *BF*_10_ = 5.28×10^6^; *t*(26) = 4.04, *p*_Holm_ < .001, *d* = .78, *BF*_10_ = 1.98 x 10^4^]. Failed STOP and IGNORE MEPs did not differ, *t*(26) = .09, *p*_Holm_ = .928, *d* = .02, BF_01_ = 8.06. Posthoc tests for Stimulation Time indicated that MEPs were higher at 150 ms than at 175 and 200 ms [respectively: *t*(26) = 2.83, p_Holm_ = .02, *d* = .55, BF_10_ = 27.73; *t*(26) = 2.44, *p*_Holm_ = .036, *d* = .47, *BF*_10_ = .97], (although Bayesian tests provide strong evidence for a difference at 175 ms and inconclusive evidence at 200 ms). Mean MEPs at the 175 and 200 ms Stimulation Times did not differ, *t*(26) = .39, *p*_Holm_ = .701, *d* = .07, *BF*_01_ = 8.85.

## Notes

### Competing Interest Statement

The authors have declared no competing interest.

